# Dissociating the effect of reward uncertainty and timing uncertainty on neural indices of reward prediction errors: A reward positivity (RewP) event-related potential (ERP) study

**DOI:** 10.1101/2021.01.06.425614

**Authors:** Alexandra M. Muir, Addison C. Eberhard, Megan S. Walker, Angus Bennion, Mikle South, Michael J. Larson

**Author notes:** Authors contributed equally to this work and are considered co-first authors (Alexandra M. Muir, Addison C. Eberhard). Correspondence to: Michael J. Larson, PhD, Brigham Young University, 244 TLRB, Provo, Utah USA 84602.

## Abstract

Accurate reward predictions include forecasting both *what* a reward will be and *when* a reward will occur. We tested how variations in the certainty of reward outcome and certainty in timing of feedback presentation modulate neural indices of reward prediction errors using the reward positivity (RewP) component of the scalp-recorded brain event-related potential (ERP). In a within-subjects design, seventy-three healthy individuals completed two versions of a cued doors task; one cued the probability of a reward outcome while the other cued the probability of a delay before feedback. Replicating previous results, RewP amplitude was larger for uncertain feedback compared to certain feedback. Additionally, RewP amplitude was differentially associated with uncertainty of presence/absence of reward, but not uncertainty of feedback timing. Findings suggest a dissociation in that RewP amplitude is modulated by reward prediction certainty but is less affected by certainty surrounding timing of feedback.

---

Reinforcement learning theory proposes that individuals monitor and adjust behavior to gain reward and avoid punishment (Holroyd & Coles, 2002; Sutton & Barto, 2015). In order to tailor behavior to achieve desired rewards, neural networks prioritize processes that accurately predict future outcomes (Schultz, 2016). When an actual outcome varies from a predicted outcome, whether positively or negatively, we experience a reward prediction error (Watabe-Uchida et al., 2017). In other words, a reward prediction error is the difference between a predicted outcome and an actual outcome (Schultz, 2016) and is thought to be directly related to subsequent behavioral adjustments to maximize future rewards (Nieuwenhuis et al., 2001; Schultz, 2016). Negative prediction errors (i.e., outcome is worse than expected) signal that behavioral change is needed (Lohse et al., 2020), whereas positive prediction errors (i.e., better than expected) indicate that no change is necessary (Lohse et al., 2020; Schultz, 2016).

The instantiation of reward prediction errors is due to a variety of neurobiological mechanisms. Prediction of future rewards occurs in networks centered around the ventral striatum (Haber, 2011) after which actual reward is compared to predicted reward in areas that include the anterior cingulate cortex, basal ganglia, and prefrontal cortex (Krigolson, 2018; Schultz, 2016; Watabe-Uchida et al., 2017). Dopamine neurons react to prediction errors in a consistently linear manner; when a reward is greater than expected, activation of dopamine neurons increases and when a reward is worse than expected, activation decreases (Schultz, 2016; Watabe-Uchida et al., 2017). When the reward is consistent with expectations, the dopamine response is decreased or absent (Schultz, 2016). As such, the response of dopamine neurons following a reward is thought to be directly related to the magnitude of the reward prediction error.

Reward prediction errors can be non-invasively, yet directly, measured using scalp-recorded brain event-related potentials (ERPs) derived from electroencephalogram (EEG) data. The reward positivity (RewP) component of the ERP is a neural reflection of reward processing, which includes predicting and evaluating rewards and experiencing reward prediction errors (Mulligan & Hajcak, 2018; Sambrook & Goslin, 2015; Schultz, 2016). The RewP is a positive going waveform that peaks at fronto-medial electrode sites approximately 200 to 300 milliseconds following the presentation of a favorable outcome or positive feedback (Proudfit, 2015). When positive feedback is absent, or if negative feedback is given, there is a negative deflection in the ERP waveform, known as the feedback-related negativity (FRN) or feedback negativity (FN). As a note, many of the papers cited in the current manuscript have investigated the FRN or the FN. However, due to recent evidence showing that the underlying ERP component is driven primarily by rewards and an absence of reward that appears as a negativity (Proudfit, 2015), the terminology RewP will be used throughout this paper. RewP amplitude is thought to be partially driven by the activity of cortical dopamine neurons (Meadows et al., 2016), and, as such, serves as a neurobiological index of reward prediction errors (Mulligan & Hajcak, 2018; Sambrook & Goslin 2015).

Along with indexing reward prediction errors, amplitude of the RewP varies based on multiple characteristics of feedback. RewP amplitude is more positive for gain trials as opposed to loss trials (in the case of the FRN, more negative for loss trials; Lohse et al., 2020; Miltner et al., 1997; Proudfit, 2015), and is greater for rewards that are larger in magnitude (Meadows et al., 2016). Uncertain outcomes also elicit greater RewP amplitude compared to certain outcomes (Hajcak et al., 2007; Pfabigan et al., 2011; Walentowska et al., 2019), but RewP amplitude may be equally effected whether the outcome is better than expected or worse than expected (Ferdinand et al., 2012; however, see Mulligan & Hajcak, 2018). Notably, higher RewP amplitude during feedback processing is associated with behavioral adjustments on subsequent trials following erroneous responses (Cohen & Ranganath, 2007; Lohse et al., 2020; Schultz, 2016), suggesting that the RewP indexes the signal between neural networks that adjusts future behavior. Taken together, most of the variation in RewP seems to be based on reward predictions and violations to said predictions (Ferdinand et al., 2012; Mulligan & Hajcak, 2018).

Although most of the variance in RewP amplitude appears to be linked to the magnitude of the reward prediction error, timing of reward has also shown an effect, with a blunted RewP amplitude following delayed feedback when compared to immediate feedback (Opitz et al., 2011a; Peterburs et al., 2016; Wang et al., 2014; Weinberg et al., 2012a; Weismüller & Bellebaum, 2016a). This idea presents the possibility that inaccuracies in the prediction of *when* an outcome will be presented may also modulate RewP amplitude. Although delayed feedback appears to result in decreased RewP amplitude compared to immediate feedback, it is possible that increasing temporal predictability of a future reward re-establishes reward prediction error as indexed by RewP amplitude. Specifically, Kimura & Kimura (2016) found that adding tones played at regular intervals leading up to a delayed reward (6,000 ms) restored RewP amplitude to its original magnitude (as seen following immediate feedback). Their results suggest that the RewP is present even during delayed feedback, but only if the feedback is able to be temporally predicted. Increasing temporal predictability of when a reward may occur may re-establish an individual’s ability to accurately evaluate action versus outcome. However, the effect of predictability of rewards and predictability of time on RewP amplitude have yet to be directly tested or dissociated.

Few studies have studied or dissociated both timing uncertainty and outcome uncertainty in the same sample. For example, startle blink response, that indexes a similar dopaminergic reward system as the RewP, appears highest during a temporally uncertain condition as compared to a certain threat, uncertain threat, and safety condition (Bennett et al., 2018). However, participants indicated on self-report measures that both the temporally uncertain conditions and outcome uncertain conditions were equally anxiety provoking, suggesting that although physiologically different, these two types of uncertainty are subjectively seen as equally distressing (Bennett et al., 2018). Heart rate also seems to increase during uncertain conditions, regardless of the temporal or reward-related nature of unpredictability (Monat et al., 1972). Although uncertain conditions seem to be driving physiological responses, Davies & Craske (2015) demonstrated that the greatest startle blink response occurred when an expected shock occurred at an expected time, compared to temporally predictable shock, temporally unpredictable shock, uncertain shock, and safe condition. Taking these studies together, the evidence surrounding the physiological response to temporal and reward unpredictability is mixed.

Individual differences can also have an impact on neurobiological indices of reward prediction such as RewP amplitude. Individuals who have greater levels of intolerance of uncertainty display greater neural response to reward prediction errors (Tanovic et al., 2018). Individuals with high trait anxiety display no difference in RewP amplitude between short and long delays in feedback presentation, while their low trait anxiety counterparts show decreased amplitude following delayed feedback (Zhang et al., 2018). Conversely, Nelson et al. (2016) found no relationship between symptoms of anxiety, depression, and worry and RewP amplitude.

Taken together, the relationship between individual differences such as intolerance of uncertainty and anxiety and RewP amplitude is unclear.

We aimed to dissociate the influence of temporal uncertainty (i.e., *when* a reward will be presented) and reward uncertainty (i.e., *what* reward will be presented) on RewP amplitude using a within-subjects design in the same experimental paradigm. First, we hypothesized that certain feedback would elicit higher RewP amplitudes than uncertain trials when cueing probability of reward and probability of time. Second, we hypothesized that RewP amplitude would be the same during timing trials that were certain, regardless of a delay, suggesting that re-establishing predictability of when a reward will occur allows for the re-establishment of reward prediction errors. Third, we hypothesized that there would be a positive correlation between measures of individual intolerance of uncertainty, anxiety, and worry with RewP amplitude, suggesting that individuals who are averse to uncertain situations display stronger neural manifestations to reward prediction errors.

## Method

All data and code used for data analyses, along with supplementary materials, have been posted to the Open Science Framework (OSF) and can be found at https://osf.io/s4m79.

### Participants and Procedures

Methods and procedures for the current study were approved by the local Institutional Review Board and all participants provided written informed consent. The initial sample consisted of 89 undergraduate students recruited for course credit. Potential participants were excluded if they had uncorrected vision difficulties, any previous head trauma that resulted in a loss of consciousness, and/or diagnosed neurological disorder (e.g., epilepsy). Participants were required to be native English speakers and be between the ages of 18 and 40. Before data analyses, seven participants were excluded for high noise levels (noise greater than 10 root mean square of the residual noise after ERP is cancelled by inverting every other trial; see Schimmel, 1967). Additionally, seven participants were excluded for not having enough trials to create a reliable signal (see Reliability and Sensitivity Analysis section below for more detail). Lastly, two participants were excluded for having outlier RewP values --outlier RewP values were identified as values outside two times the inter-quartile range (Gareth, Witten, Hastie, & Tibshirani, 2020). The final sample size included 73 healthy young adult participants (36 male (49.3%); *M(SD)*_age_ = 20.7(2.0)).

Participants completed one 75-minute laboratory session wherein they provided written consent followed by a battery of questionnaires that included a standard demographic questionnaire, the Intolerance of Uncertainty-Short Form (IU-12), the Penn State Worry Questionnaire (PSWQ), the State-Trait Anxiety Inventory-Trait Subscale (STAI-Trait), and the Mood and Anxiety Symptom Questionnaire (MASQ). A trained research assistant subsequently administered the Test of Premorbid Functioning from Advanced Clinical Solutions (TOPF) as a measure of estimated intelligence. All questionnaires are described in the following section. Upon completion, participants were fitted with an electroencephalogram cap after which they completed two modified versions of the doors task (Proudfit et al., 2015).

A schematic representation of the two experimental tasks is provided in Figure 1. Both tasks were identical except for the meaning of the shape cues presented before each trial, as described in the following paragraph. The order of tasks was counterbalanced between participants. At the start of the task, a white shape (either square, triangle, or diamond) on a black background was displayed for 2000 ms, after which a fixation cross was displayed for 500 ms. Then, two identical doors were shown, and the participant was instructed to pick either the left or right door. After clicking a mouse button to pick a door, an additional fixation cross was shown for jittered intervals of either 500 to 1000 ms for the short fixation crosses and 3500 to 4000 ms in the reward-cued task and 4500 to 5000 ms in the timing-cued task for the long fixation crosses. Participants were then shown either a green arrow pointing upwards (signifying a win of 20 cents) or a red arrow pointing downwards (signifying a loss of 10 cents). Rewards were twice as large as losses due to evidence that losses are subjectively twice as salient as gains (Tversky & Kahneman, 1992). For each task, participants completed 160 trials, with feedback being fixed at 50% gains and 50% losses, although participants were not made aware of this during the task to ensure that a feedback ERP was elicited. In both tasks, there were 8 trial types (with short and long referring to the delay of feedback): certain gain short, certain gain long, certain loss short, certain loss long, uncertain gain short, uncertain gain long, uncertain loss short, and uncertain loss long. 20 trials of each type were displayed in a random order.

**Figure 1:**
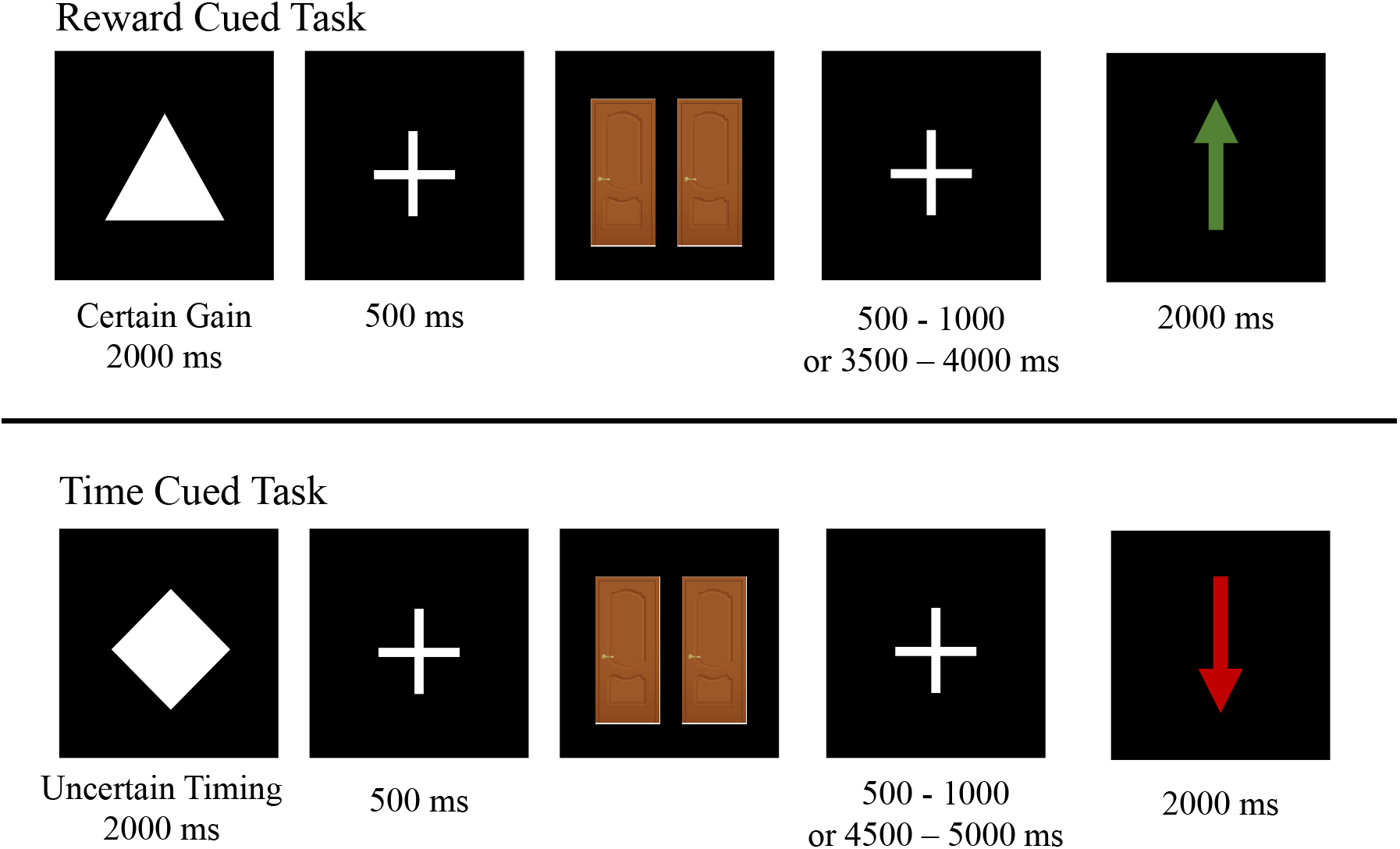
Schematic representation of experimental tasks

The only difference between the two tasks was the meaning of the shape cues that were shown before the doors. In the reward-based task, if a square was seen before the doors appeared the participant knew that there was a 100% chance of winning money (certain gain) while a triangle meant 100% chance of losing money (certain loss). The last cue, a diamond, meant that the feedback was dependent on the participant’s response (uncertain), although feedback was actually randomized. In the timing-based task, the cues referenced the length of the fixation cross preceding the presentation of feedback rather than the feedback itself. A square meant that feedback would be presented quickly (between 500 and 1000 ms), while a triangle meant that feedback would be delayed (3500 to 4000 ms for reward-cued task or 4500 to 5000 ms for time-cued task; difference was due to a design error). A diamond meant that the length of time spent waiting between door choice and feedback receipt was unknown to the individual (i.e., uncertain). Participants were told that their reward still depended on what door they picked, regardless of the delay of feedback. All other aspects of the tasks were identical. Before both tasks, the participant first completed a six-trial practice task to ensure they fully understood the meaning of the shape cues. After finishing both tasks participants were compensated for their time either through course credit or monetary payment ($20 total for completing the entire study).

### Reliability and Sensitivity Analyses

To determine the number of trials needed to create a reliable signal, dependability estimates were determined using the ERP Reliability Analysis Toolbox v0.3.2 (Clayson & Miller, 2017). All dependability estimates are presented in Table 1. Dependability estimates for the time-cued task were > 0.83; dependability estimates for the reward-cued task were > 0.70. For the reward-based task, greater than 22 trials in each category was needed to achieve adequate reliability (defined as α = 0.6; see Table 1 for trial cutoffs per trial type). For the timing-based task, a minimum number of 8 useable trials in any of the trial types were needed to reach the 0.6 cutoff. Reliability cut-offs are in-line with previous estimates for the RewP and FN (Ethridge & Weinberg, 2018).

**Table 1:** ERP dependability and error estimates

| Trial Type | Dependability | Mean(SD) Trials | Trial Range | Minimum Trials | Noise Mean(SD) |
| --- | --- | --- | --- | --- | --- |
| Reward-Cue |  |  |  |  |  |
| Certain Gain | 0.75 | 33.0(5.2) | 19, 40 | 17 | 1.4(0.6) |
| Certain Loss | 0.70 | 34.1(4.6) | 22, 40 | 22 | 1.4(0.5) |
| Uncertain Gain | 0.93 | 34.2(5.2) | 20, 40 | 4 | 1.4(0.7) |
| Uncertain Loss | 0.91 | 33.7(5.5) | 19, 40 | 5 | 1.4(0.5) |
| Time-Cue |  |  |  |  |  |
| Certain Gain | 0.85 | 24.7(4.3) | 10, 29 | 7 | 1.6(0.6) |
| Certain Loss | 0.83 | 25.7(4.3) | 9, 30 | 8 | 1.6(0.9) |
| Uncertain Gain | 0.90 | 34.2(6.2) | 11, 40 | 6 | 1.4(0.6) |
| Uncertain Loss | 0.86 | 34.4(5.4) | 15, 40 | 8 | 1.4(0.6) |

To ensure that the current study was adequately powered to the effects of interest (Larson & Carbine, 2017), a sensitivity analysis was performed in G*Power (v.3.1). For the total sample size of 73, with an alpha of 0.05, power of 0.80, eight repeated measures, and a 0.4 correlation between repeated measures, we were powered to detect an *f* of 0.12. This indicates the current study was adequately powered to detect small to medium effects present in the data.

### Questionnaires

All means, standard deviations, and Cronbach’s alphas for the questionnaire measures in the current sample are presented in Table 2. The IU-12 is a 12-statement questionnaire that aims to quantify an individual’s intolerance of uncertainty along with their predisposition towards disliking uncertain future events (Alexander & Brown, 2011). Participants respond to statements such as “Unforeseen events upset me greatly” and “I must get away from all uncertain situations” on a scale ranging from “not at all characteristic of me” (+1) to “entirely characteristic of me” (+5; Oglesby et al., 2017). Item responses are summed together to create a composite score, with a greater score indicating greater intolerance towards uncertainty. Total scores on the IU-12 can range from 12 to 60. Along with measuring overall displeasure towards uncertainty, the intolerance of uncertainty scale can also measure the subcomponents of prospective anxiety and behavioral anxiety.

**Table 2:** Questionnaire information

|  | Mean | SD | Range | Cronbach's Alpha |
| --- | --- | --- | --- | --- |
| IU-12 | 29.59 | 7.9 | 14, 54 | 0.87 |
| IU-Prospective | 19.60 | 4.9 | 9, 33 | 0.82 |
| IU-Inhibitory | 10.00 | 3.9 | 5, 22 | 0.82 |
| PSWQ | 42.29 | 7.36 | 29, 62 | 0.95 |
| STAI-State | 44.24 | 4.39 | 34, 55 | 0.93 |
| MASQ-Anxious Arousal | 21.33 | 3.94 | 17, 35 | 0.67 |

Along with the IU-12, three difference anxiety scales were administered to participant, namely the PSWQ, the STAI-Trait, and the MASQ-AA. All three questionnaires were used for analyses to characterize a broader picture of anxiety symptomology in participants. First, the PSWQ aims to quantify levels of worry (Meyer et al., 1990). Sixteen items such as “my worries overwhelm me” or “I do not tend to worry about things” are ranked using a Likert-type scale ranging from “not at all typical of me” (+1) to “very typical of me” (+5). Items were reverse scored as needed and summed for a total score that ranges between 16 to 80 points. A higher score indicates higher levels of trait worry. The PSWQ generally has a high internal consistency with a Cronbach’s alpha of 0.95 (Meyer et al., 1990).

The STAI-Trait form Y-2 also quantifies anxiety symptoms (Speilberger et al., 1970). For the state subscale, twenty items such as “I feel calm” were rated on a four-point Likert-type scale ranging from “not at all” (+1) to “very much so” (+4). Items were reverse scored as needed, and then summed for a total score. The STAI-state subscale can range from a score of 20 to 80. Overall, the STAI has good internal consistency, with Cronbach’s alpha above 0.7 (Bergua et al., 2012; Speilberger et al., 1970).

Lastly, the MASQ aims to quantify both anxiety and depressive symptomology in an individual (Watson et al., 1995). The full MASQ is 90 items, however, in the current study we focused in on the MASQ Anxious Arousal (MASQ-AA) subscale. As such, participants rated thirty-eight statements such as “hands were shaky” or “felt cheerful” on a five-point scale ranging from “not at all” (+1) to extremely (+5). Seventeen out of the thirty-eight items directly related to the MASQ-AA subscale, and were summed for a total MASQ-AA score. Scores for the anxious arousal subscale range from 17 to 85. The MASQ generally shows good psychometric properties and provides strong internal validity (Lin et al., 2014).

The Test of Premorbid Functioning (TOPF) from the Advanced Clinical Solutions was used as an intelligence estimate. To complete the TOPF participants read a list of 70 irregular words during which the administrator of the test determined if the word had been pronounced correctly. At the end, correctly pronounced words were summed, and standardized scores based on age were determined using a standardized booklet (Psychological Corporation, 2009). The TOPF displays similar scores as other tests of pre-morbid functioning (Kirton et al., 2020), although some have argued that the TOPF just reflects English word knowledge rather than the intelligence construct itself (Shura et al., 2020) or may underpredict premorbid intelligence in certain populations (Berg et al., 2016; Joseph et al., 2019). The TOPF was used in the current study to estimate the intelligence level of study participants. Internal reliability scores for the TOPF were between 0.89 and 0.95 and the test-retest correlations were between 0.89-0.95 (Chu et al., 2012). The TOPF is also highly correlated with other tests designed to measure verbal functioning (Chu et al., 2012) The current sample displayed a mean intelligence estimate of 107.3 (*SD* = 10.9).

### Electroencephalogram recording and reduction

All EEG data were recorded through 128 passive Ag/AgCl electrodes on a Hydrocel geodesic sensor net from Electrical Geodesics, Inc. Net-Amps 300 system with a nominal gain and online bandpass filter from .01 to 100 Hz. Data were referenced to the vertex electrode Cz during data collection and digitized continuously at 250 Hz with a 16-bit analog-to-digital converter. Impedances were kept under 50 kΩ per the manufacture’s recommendation. After finishing data collection, data were digitally offline high-pass filtered with a 0.05 Hz filter and digitally low-pass filtered at 15 Hz in NetStation (v. 5.3.0.1). Data were segmented from 200 ms before reward stimulus onset until 800 ms after. The following segments were pulled for both the reward and timing tasks: certain short, certain long, uncertain short, uncertain long, certain reward, certain loss, uncertain reward, uncertain loss. Artifacts were then corrected using independent components analysis (ICA) in the ERP PCA toolkit in Matlab (Dien, 2010). If any ICA component correlated with two blink templates (one template from the ERP PCA toolkit and one made previously by the authors) at a rate of 0.9 or higher, that component was removed from the data. Additionally, if the fast average amplitude of a channel was greater than 100 microvolts, differential average amplitude was less than 50 microvolts, or the eye-channel amplitudes surpassed 70 microvolts, that particular channel was defined as unacceptable and the six closest nearest-neighbor electrodes were used to interpolate and replace the electrode. Following artifact correction, all data were re-referenced to the average reference and baseline adjusted from −200 to 0 ms before feedback presentation.

Following baseline adjustment, as decided *a* priori, we used the collapsed localizer approach by collapsing across all trial types to view an overall grand-averaged waveform to select the RewP/FN window (Luck & Gaspelin, 2017). Feedback ERP data were then extracted as the mean amplitude from 170 to 300 ms following feedback onset. Mean amplitude was chosen due to evidence suggesting that mean amplitude is more reliable than other ERP peak measures (Clayson et al., 2013; Luck, 2005). Event-related potential data values were extracted using R (v 1.1.463; R Core Team, 2020) and averaged over four fronto-central electrodes (6 [FCz], 7, 106, 129 [Cz]). A region of interest average was used based on data showing improved reliability when using a sensor average over single electrodes (Baldwin et al., 2015; Clayson, 2020). Means and standard deviations for the RewP by trial type and task are presented in Table 3.

**Table 3:** ERP and response time values by trial type and task

|  | Reward Task |  |  | Time Task |  |  |
| --- | --- | --- | --- | --- | --- | --- |
| RewP Amplitude (μV) | Mean | SD | Range | Mean | SD | Range |
| Certain Gain | 1.8 | 1.6 | -2.5, 6.4 | 4.2 | 2.7 | -1.3, 12.6 |
| Certain Loss | 1.5 | 1.4 | -1.4, 5.5 | 3.3 | 2.5 | -3.3, 11.5 |
| Uncertain Gain | 4.5 | 3.5 | -0.8, 18.4 | 4.3 | 2.9 | -0.3, 13.1 |
| Uncertain Loss | 3.8 | 3.2 | -2.2, 16.4 | 3.6 | 2.5 | -1.6, 10.0 |
| Response Time (ms) | Median | SD | Range | Median | SD | Range |
| Overall RT | 563.5 | 177.4 | 288.5, 1156.5 | 606.0 | 430.4 | 266.5, 2641.5 |
| Certain Gain | 463.5 | 187.6 | 268.0, 1337.5 | 580.5 | 680.0 | 240, 5418.5 |
| Certain Loss | 525.0 | 226.3 | 246.5, 1718.5 | 587.0 | 465.6 | 270.5, 2504.5 |
| Uncertain Gain | 608.0 | 290.8 | 323.5, 1884.5 | 623.5 | 426.9 | 258.5, 2843.5 |
| Uncertain Loss | 585.0 | 271.8 | 287.0, 1605.5 | 588.0 | 433.9 | 298, 2976.5 |

### Data Analysis

All ANOVAs described below were conducted using the “ez” package in R (Lawrence, 2016). For behavioral data, median response time was calculated across all trials in both tasks. A 2-task (reward, time) x 2-certainty (certain, uncertain) x 2-outome (gain, loss) ANOVA was conducted on median response time to determine if response time varied based on certainty or outcome by task.

Two repeated-measures analyses of variance (ANOVA) were subsequently conducted to test the relationship between reward and timing uncertainty ERP amplitude. The first ANOVA aimed to parse apart certainty differences in reward outcome using a 2-task (reward, time) x 2-certainty (certain, uncertain) x 2-outcome (gain, loss) ANOVA. The second ANOVA aimed to parse apart certainty differences in timing outcome using a 2-task (reward, time) x 2-certainty (certain, uncertain) x 2-timing (short, long) ANOVA. Significant interaction effects were decomposed using *t*-tests or additional ANOVAs (in the case of three-way interactions). Generalized eta squared is presented as a measure of effect size for all ANOVAs.

As noted above, outlier RewP values were determined as participants whose RewP amplitude values were outside two times the inter-quartile range (Gareth, Witten, Hastie, & Tibshirani, 2020). Outliers were examined for each ANOVA. The first ANOVA on reward certainty had two participants excluded (final sample size: 73; 36 male (49.3%); *M(SD)*_age_ = 20.7 (2.0)) and the second ANOVA on timing uncertainty had four participants excluded (final sample size: 71; 34 male (40.9%); *M(SD)*_age_ = 20.7 (2.0)).

Exploratory correlations between RewP amplitude and IU-12, IU-prospective anxiety, IU-behavioral anxiety, PSWQ, MASQ-AA and STAI-State were conducted to see how neural indices of reward processing related to self-reported intolerance of uncertainty and various dimensions of anxiety symptomology. Since certain and uncertain trials were not significantly different in the time task, only RewP amplitudes from the reward-based task are used in the correlational analyses. Both certain and uncertain trial RewP amplitude from the reward-based task were separately correlated with total scores on the IU-12, PSWQ, and MASQ-AA and subscale scores of IU-prospective anxiety and IU-behavioral anxiety.

## Results

Median response times as a function of task and trial type are shown in Table 2. A 2-task (reward, time) by 2-certainty (certain, uncertain) by 2-outcome (gain, loss) ANOVA showed a no main effect of certainty (*F*(1,72) = 4.0, *p* = .05, η^2^ = 0.002), but a main effect of task, (*F*(1,72) = 7.4, *p* = .01, η^2^ = .03) and a task by certainty interaction (*F*(1,72) = 12.1, *p* < .01, η^2^ = 0.01). Response times were slower for uncertain trials compared to certain trials on the reward-cued task (*t*(145) = −4.6, p < .01, *d*_z_ = −0.38) but not on the time-cued task (*t*(145) = .82, *p* = .41, *d*_*z*_ = 0.05). The time-cued task had longer RT compared to the reward-cued task for both certain (*t*(145) = −4.3, *p* < .01, *d*_*z*_ = −0.42) and uncertain (*t*(145) = −2.2, *p* = .03, *d*_*z*_ = −0.21) trials.

RewP ERP component waveforms for each task are displayed in Figure 2. A 2-task (reward, time) x 2-certainty (certain, uncertain) x 2-outcome (gain, loss) ANOVA showed a main effect of task (*F*(1,72) = 20.1, *p* < .01, η^2^ = 0.04), a main effect of certainty (*F*(1,72) = 75.6, *p* < .01, η^2^ = 0.07), and a main effect of outcome (*F*(1,72) = 34.8, *p* < .01, η^2^ = .02). The timing task showed larger amplitudes than the reward task (*M(SD)*_*Time*_ = 3.7 μV (2.4), *M(SD)*_*Reward*_ = 2.8 μV (2.6)), uncertain trials showed greater RewP amplitude than certain trials (*M(SD)*_*Uncertain*_ = 3.9 μV (2.7), *M(SD)*_*Certain*_ = 2.6 μV (2.2)), and gain trials had greater amplitude than loss trials, as expected (*M(SD)*_*Gain*_ = 3.5 μV (2.6), *M(SD)*_*Loss*_ = 2.9 μV (2.4)). There was no certainty by outcome interaction (*F*(1,72) = .04, *p* = .84, η^2^ = 0.002) nor a significant task by outcome interaction (*F*(1,72) = 3.9, *p* = .05, η^2^ < 0.01). There was, however, a significant task by certainty interaction (*F*(1,72) = 58.6, *p* < .01, η^2^ = 0.05). Importantly, however, all of these main effects and interactions were superseded by a significant task by certainty by outcome three-way interaction (*F*(1,72) = 4.5, *p* = .04, η^2^ = 0.001).

**Figure 2:**
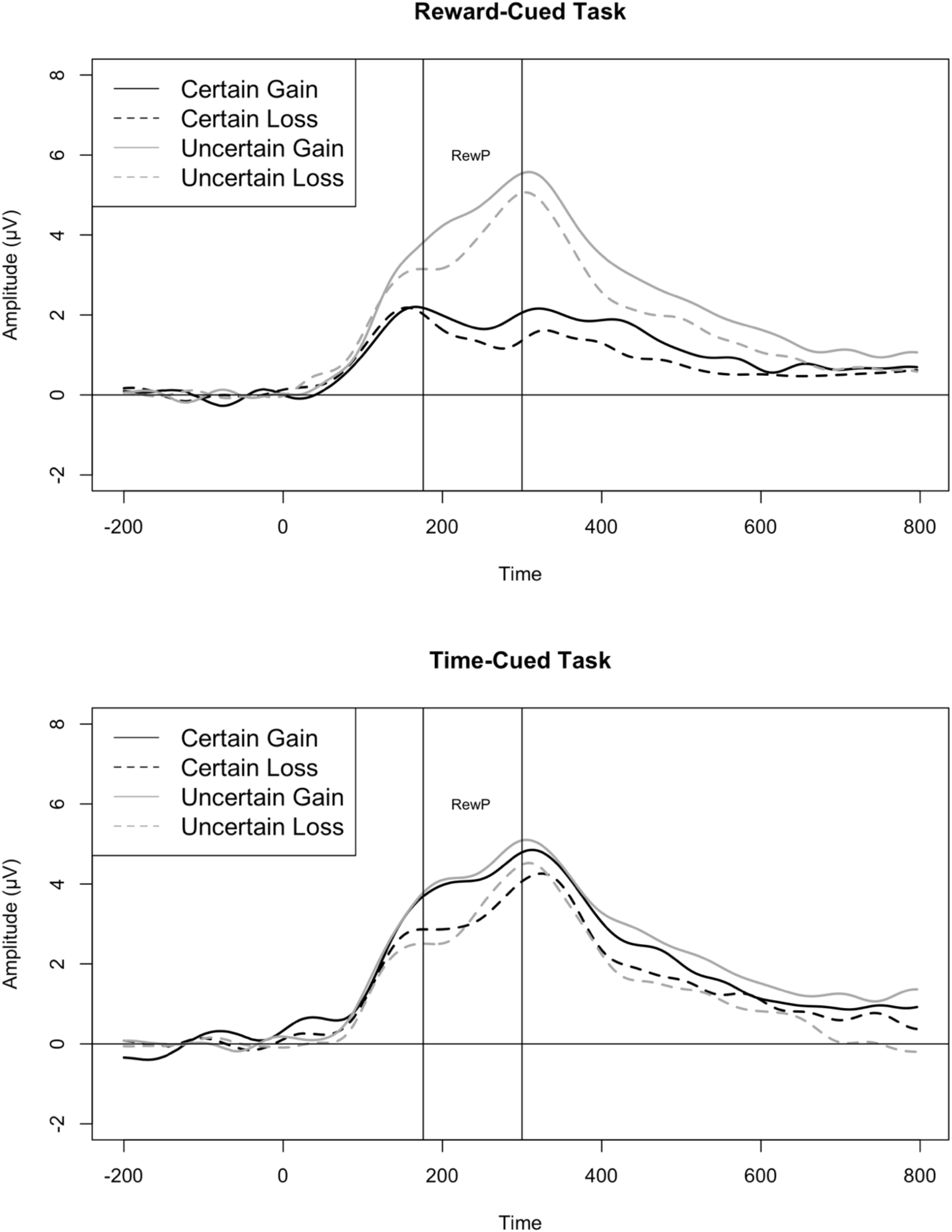
Grand-averaged waveforms for the RewP for both the reward-cued and time-cued tasks averaged over four fronto-central electrodes.

Figure 3 shows visual representation of the means to decompose the three-way interaction. Specifically, to understand the three-way interaction, follow-up ANOVAs were conducted, with a 2-task (reward, time) by 2-certainty (certain, uncertain) being conducted for both gain trials and loss trials separately. For gain trials only, there was a significant main effect of certainty (*F*(1,72) = 64.4, *p* < .01, η^2^ = 0.18) and task (*F*(1,72) = 21.4, *p* < .01, η^2^ = 0.13), along with a significant certainty by task interaction (*F*(1,72) = 53.3, *p* < .01, η^2^ = 0.15). When decomposing the significant certainty by task interaction for gain trials, uncertain trials resulted in larger RewP amplitudes at time of feedback compared to certain trials for the reward-cued task (*t*(72) = −9.1, *p* < .01, *d*_*z*_ = −0.95) but not the time-cued task (*t*(72) = −0.67, *p* = .51, *d*_*z*_ = −0.05). Looking between the two tasks, certain trials (with eventual outcome of gain) elicited higher RewP amplitudes during the timing-cued task when compared to the reward-cued task (*t*(72) = −8.5, *p* < .01, *d*_*z*_ = −1.1), but there were no differences in RewP amplitude between the reward-cued and time-cued task for uncertain trials (*t*(72) = 0.32, p = .75, d_z_ = .04). For the loss trials, there was a main effect of certainty (*F*(1,72) = 50.3, *p* < .01, η^2^ = 0.19), task (*F*(1,72) = 12.1, *p* < .01, η^2^ = 0.07), and a significant task by certainty interaction (*F*(1,72) = 38.6, *p* < .01, η^2^ = 0.11). Uncertain trials that resulted in a loss displayed higher RewP amplitude during both the reward-(*t*(72) = −7.5, *p* < .01, *d*_*z*_ = −0.05) and timing-cued tasks (*t*(72) = −2.6, *p* = .01, *d*_*z*_ = −0.16). Similar to the gain-only ANOVA, certain trials elicited higher RewP amplitudes during the timing-cued task compared to the reward-cued task (*t*(72) = −6.4, *p* < .01, *d*_*z*_ = −0.84), but there were no differences observed between the two tasks for the uncertain trials (*t*(72) = 0.64, *p* = .53, *d*_*z*_ = .06).

**Figure 3:**
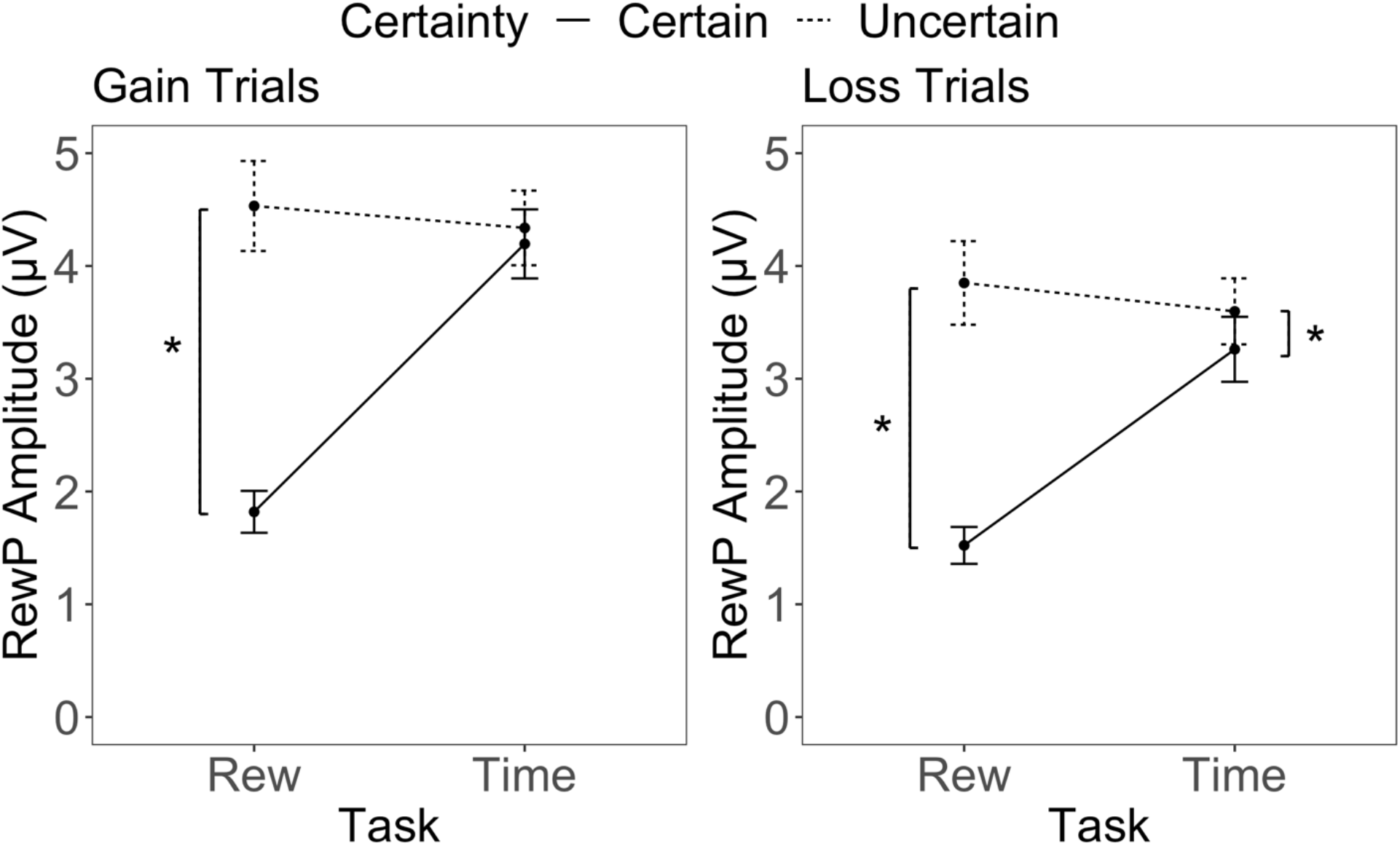
Interaction plots of the three-way interaction. Task by certainty interaction split by outcome type (gain, loss) Significant differences are marked with an *.

**Figure 4:**
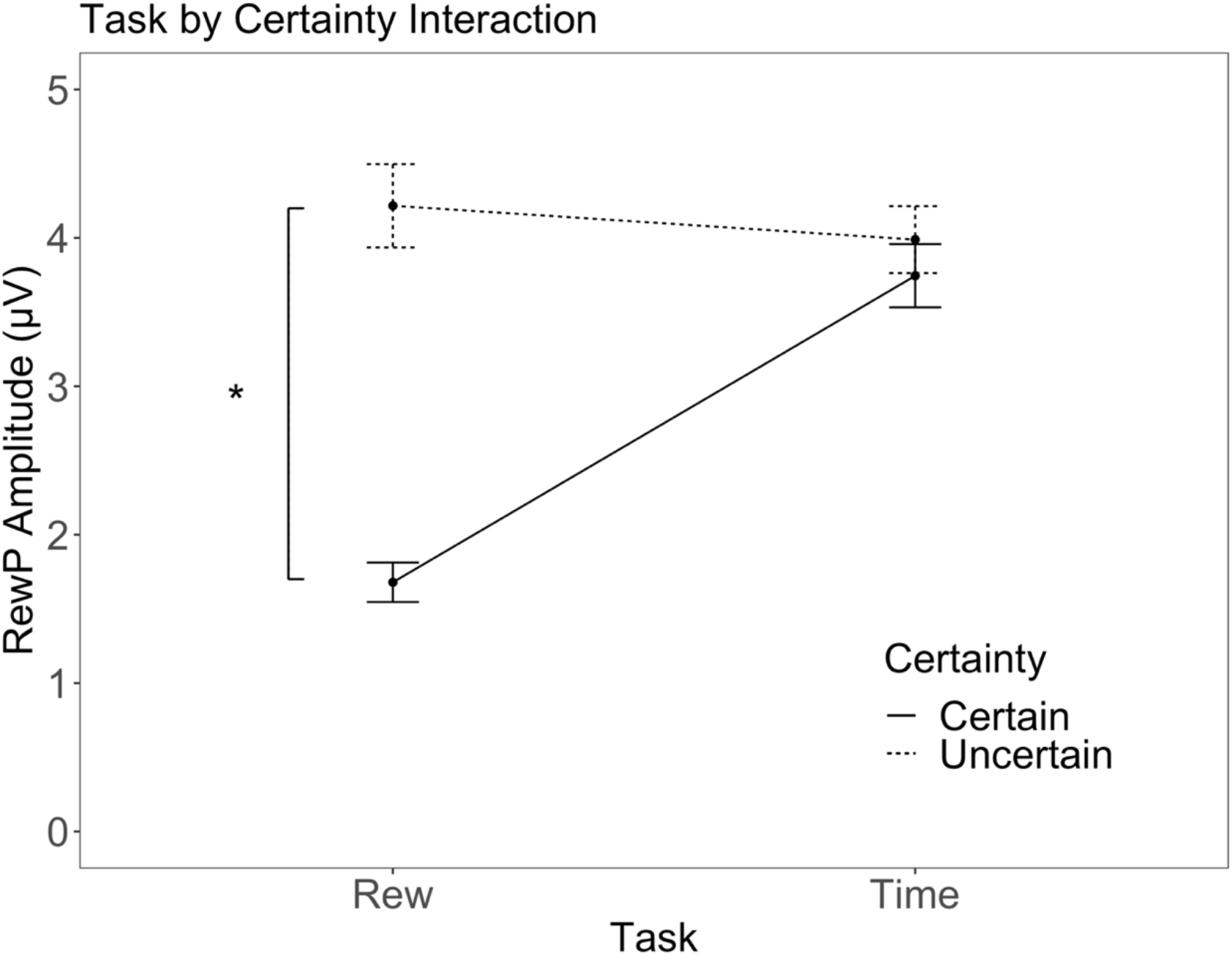
Task by certainty interaction for the secondary ANOVA examining task by certainty by fixation cross.

A 2-task (reward, time) x 2-certainty (certain, uncertain) x 2-length (long, short) ANOVA displayed a similar main effect of task (*F*(1,70) = 19.3, *p* < .01, η^2^ = 0.08) and a main effect of certainty (*F*(1,70) = 71.4, *p* < .01, η^2^ = 0.14). Again, the time task and uncertain trials elicited greater RewP amplitudes on average. There was no main effect of fixation cross length (*F*(1,70) = 2.4, *p* = .12, η^2^ = 0.008). There was also no three-way interaction (*F*(1,70) = 0.32, *p* = .57, η^2^ < 0.001), nor a certainty by length interaction (*F*(1,70) = 3.3, *p* = .07, η^2^ = 0.003). There was, however, a task by certainty interaction (*F*(1,70) = 54.2, *p* < .01, η^2^ = 0.10) and a task by length interaction (*F*(1,70) = 16.7, *p* < .01, η^2^ = 0.01). These interactions are depicted in Figure 5 and Figure 6. Uncertain trials were more positive than certain trials just for the reward-based task (*t*(141)_Reward_ = −11.2, *p* < .01, *d*_*z*_ = −0.87; *t*(141)_Time_ = −1.9, *p* = .05, *d*_*z*_ = −0.10). For certain trials, the time task elicited higher RewP amplitudes compared to the reward task (*t*(141) = −9.9, *p* < .01, *d*_*z*_ = −0.94) but no differences were present for the uncertain trials (*t*(141) = 0.55, *p* = .58, *d*_*z*_ = 0.04). For the task by length interaction, the long fixation cross elicited a more positive RewP compared to the short fixation cross, but only for the reward-cued task (*t*(141)_Reward_ = 4.1, *p* < .001, *d*_*z*_ = 0.24; *t*(141)_Time_ = −0.48, *p* = .64, *d*_*z*_ = −0.03). The time-cued task elicited greater RewP amplitude compared to the reward-cued task for both the long fixation cross trials (*t*(141) = −2.6, *p* = .01, *d*_*z*_ = −0.21) and the short fixation cross trials (*t*(141) = −5.9, *p* < .001, *d*_*z*_ = −0.51).

**Figure 5:**
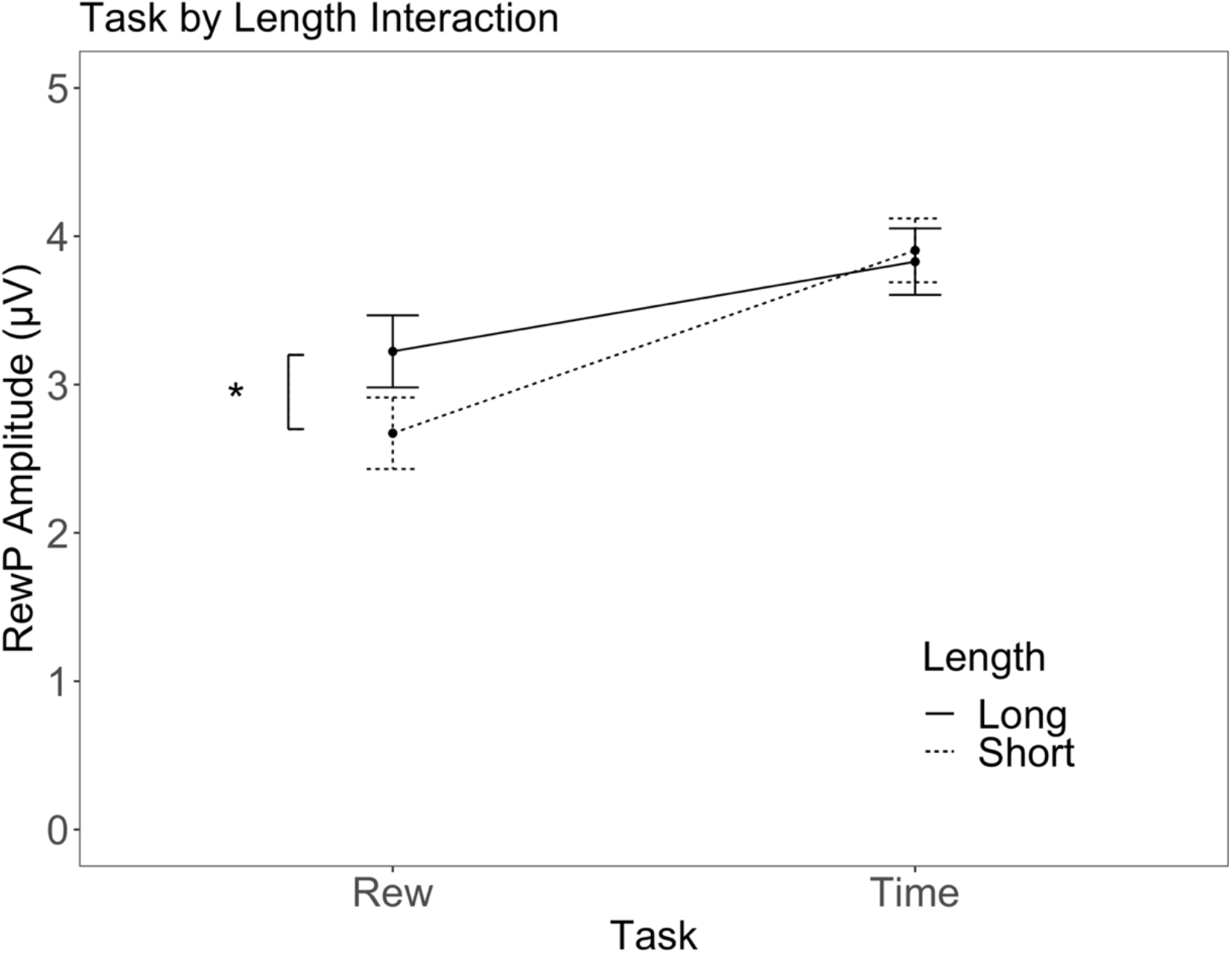
Task by fixation cross length interaction for secondary ANOVA.

Correlations between questionnaires and RewP amplitudes (only for the reward-cued based task) are presented in Table 5 while scatterplots are displayed in the supplementary materials, namely Figures S1 through S6. All correlations between individual differences measures and RewP amplitude were non-significant (*r*s < 0.12, *p*s > .41).

**Table 5:** Correlations of RewP Amplitude with Questionnaire Scores

|  | IU-12 |  | IU-Prospective |  | IU-Inhibitory |  | MASQ-AA |  | PSWQ |  | STAI-State |  |
| --- | --- | --- | --- | --- | --- | --- | --- | --- | --- | --- | --- | --- |
|  | r | p | r | p | r | p | r | p | r | p | r | p |
| Certain Gain | 0.01 | 0.92 | 0.00 | 0.98 | 0.02 | 0.85 | -0.02 | 0.84 | -0.07 | 0.47 | 0.00 | 0.99 |
| Certain Loss | 0.04 | 0.73 | 0.04 | 0.75 | 0.04 | 0.77 | -0.06 | 0.60 | -0.08 | 0.51 | -0.10 | 0.41 |
| Uncertain Gain | 0.07 | 0.54 | 0.12 | 0.33 | 0.00 | 0.99 | 0.00 | 0.97 | 0.01 | 0.93 | 0.02 | 0.92 |
| Uncertain Loss | 0.03 | 0.81 | 0.06 | 0.64 | -0.01 | 0.91 | 0.02 | 0.85 | 0.00 | 0.99 | 0.03 | 0.80 |

## Discussion

The current study demonstrated a dissociation between neurophysiological responses to uncertainty of reward and uncertainty of timing using a within-subjects design sensitive to small-to-medium effects in a sample of healthy undergraduate students. Results suggest that the RewP component primarily indexes reward-specific prediction errors rather than errors in prediction of timing. Additionally, the current study provides evidence for increased motivational salience of rewards compared to the timing of reward or non-reward presentation (cf., Pegg & Kujawa, 2020). As expected, we found that larger RewP amplitudes were present on uncertain reward trials compared to certain reward trials, confirming previous studies that have shown the same or similar effect (Hajcak et al., 2007; Pfabigan et al., 2011; Walentowska et al., 2019). However, contrary to our hypothesis, RewP amplitude did not significantly differ between certain and uncertain trials on the timing cued task where participants were cued to the timing of reward presentation but not the presence/absence of reward. A similar pattern of results appeared for the behavioral data, with response times being slower for uncertain trials compared to certain trials, but only for the reward-cued task and not for the time-cued task.

There are many neurobiological mechanisms involved in reward anticipation and processing, including the ventral striatum, amygdala, medial prefrontal cortex, caudate, and orbitofrontal cortex (Carlson et al., 2011). The ventromedial prefrontal cortex is thought to be involved in the anticipation of rewards while dopaminergic midbrain regions evaluate reward prediction errors and signal the need for an update in future behavior (Iigaya et al., 2020). Separately, temporal processing is localized to the ventral striatum (Iigaya et al., 2020). Although all of these neural sources contribute to the RewP (Carlson et al., 2011; Foti et al., 2011), it is likely that in healthy undergraduate students, the anticipation and salience of the reward itself outweighs the anticipation of when the reward will occur. As such, neural activity may be centered around the prediction and anticipation of reward valence rather than reward timing.

The difference in RewP amplitude for certain and uncertain trials in the reward task and lack thereof in the timing task provides a baseline understanding of the dissociation of reward and temporal processing in healthy adults, while providing a foundation for investigating the same processes in those with psychopathology. Aberrant reward processing is present in a number of clinical mental health conditions, including depression (Clayson et al., 2020; Keren et al., 2018; Nusslock & Alloy, 2017), autism (specifically of social rewards; Stavropoulos & Carver, 2014), attention-deficit/hyperactivity disorder (Kallen et al., 2020), and anxiety (Gu et al., 2010; Stavropoulos & Carver, 2014). It is possible that those with psychopathology do not demonstrate the same differential pattern between certain and uncertain reward/timing trials as non-clinical individuals. For example, current hypotheses surrounding reward anticipation in anxiety state that although temporal ambiguity does not seem to affect non-clinical participants, it is possible that ambiguity in the timing of a reward may leave room for negative interpretation bias and increased distress in individuals with psychopathology (Carleton et al., 2012). Similarly, children and adolescents may show increased anticipation under certain or uncertain situations for reward presence or reward timing compared to adults (Kujawa et al., 2018; Moser et al., 2018). Future iterations of the current study could investigate if the same lack of differentiation between uncertain and certain timing trials is present in those with psychopathology and across development.

The salience of reward valence is also evidenced by a lack of difference in RewP amplitude between short and delayed trials in the timing-cued task. Unlike previous studies, where delayed feedback resulted in a blunted RewP amplitude (Arbel et al., 2017; Opitz et al., 2011b; Qu et al., 2013; Weinberg et al., 2012b; Weismüller & Bellebaum, 2016b) RewP amplitude on uncertain delayed trials was not significantly different than uncertain short trials. Although our second hypothesis stated that reestablishing predictability of reward timing would restore RewP amplitude, RewP amplitude was not blunted following delayed feedback in the first place, leaving nothing to restore. The current study also found that during the reward-cued task, trials with immediate feedback actually had smaller RewP amplitude compared to delayed feedback. The presentation of immediate feedback may have been too quick for the participant to develop a significant anticipation (i.e. expectation) of future reward. However, it is also possible, as noted above, that the salience of the reward simply outweighs the timing ambiguity between the cue and reward presentation. In contrast to a recent study of attention bias modification that showed no changes in RewP amplitude despite changes in attention to other components (Sylvain et al., 2020), the blunted RewP amplitude for short trials suggests attention and time to attend to reward may influence RewP amplitude outcomes.

Consistent with previous studies, no individual difference measure of anxiety nor intolerance of uncertainty correlated with RewP amplitude on the uncertain trials (Nelson et al., 2016). Specifically, Nelson et al., (2016) found no relationship between measures of worry, depressive symptomology and stress with RewP amplitude (Nelson et al., 2016). However, our results are inconsistent with Nelson et al. (2016) when looking at intolerance of uncertainty, as they did find that inhibitory intolerance of uncertainty was associated with blunted RewP amplitude while prospective intolerance of uncertainty was associated with increased RewP amplitude. The current study found no correlation between overall intolerance of uncertainty and RewP amplitude and no correlation between the subscales of prospective and inhibitory anxiety and RewP amplitude. Recent work shows that intolerance of uncertainty is related to error-related neural processes and depressive symptoms (Ruchensky et al., 2020). Given that intolerance of uncertainty tends to be heightened in people with internalizing psychopathology who may attend more to threat during unpredictable conditions (Correa et al., 2019; Imburgio & MacNamara, 2019), the current healthy undergraduate sample may simply not have sufficient range on the intolerance of uncertainty scale to significantly correlate with reward processing outcomes.

Our results should be understood in the context of the study limitations and strengths. The time window of delayed feedback was shorter than other studies examining the effect of timing delays on RewP amplitude. As such, it is possible that the delayed condition in the current paradigm (3500 ms −5000 ms) could be considered a “medium” delay rather than a “long” delay (6000 or more ms; Peterburs et al., 2016; Weinberg et al., 2012), and could contribute to the difference in results between the current study and previous studies. Additionally, due to an error in task programming error, the duration of the delayed condition was different across the reward and timing tasks. The reward task had a jittered delay of 3500 ms to 4000 ms while the timing task had a jittered delay of 4500 ms to 5000 ms. Although the difference of approximately 1000 ms is not likely to cause substantial differences in results, it is possible that this lack of standardization added noise into the analyses presented in the paper. With that said, we used a relatively high-powered, within-subjects design and sensitivity analyses show that using the current sample size and design we had the ability to detect small and medium effects present in the data. Additionally, we were able to test the effect of reward and temporal predictability on RewP amplitude in the same paradigm in a within-subjects design, allowing us to make more consistent comparisons between reward and temporal uncertainty.

In conclusion, the results of this study add to the current body of literature in discussing the underlying generators of the RewP by directly comparing effects of reward and timing uncertainty on RewP amplitude. The current results confirm that uncertain trials elicit greater RewP amplitude, when compared to certain trials (Hajcak et al., 2007; Pfabigan et al., 2011; Walentowska et al., 2019; Xu et al., 2011), which supports theoretical arguments that the RewP indexes a reward prediction error. These results also add, however, that this effect is driven by uncertainty specifically surrounding the reward and is unchanged by uncertainty surrounding the timing of feedback presentation. This evidence dissociation in RewP amplitude for certain and uncertain trials in relation to rewards and not in relation to timing provides a baseline understanding of reward and temporal processing in healthy adults and should be further probed in individuals with psychopathology.

## Author Note

This research was supported by funds from the Brigham Young University College of Family, Home, and Social Sciences.

There was no conflict of interest while conducting this study.

