## Supplemental Material for "Dissociating the effect of reward uncertainty and timing uncertainty on neural indices of reward prediction errors: A reward positivity (RewP) event-related potential (ERP) study"

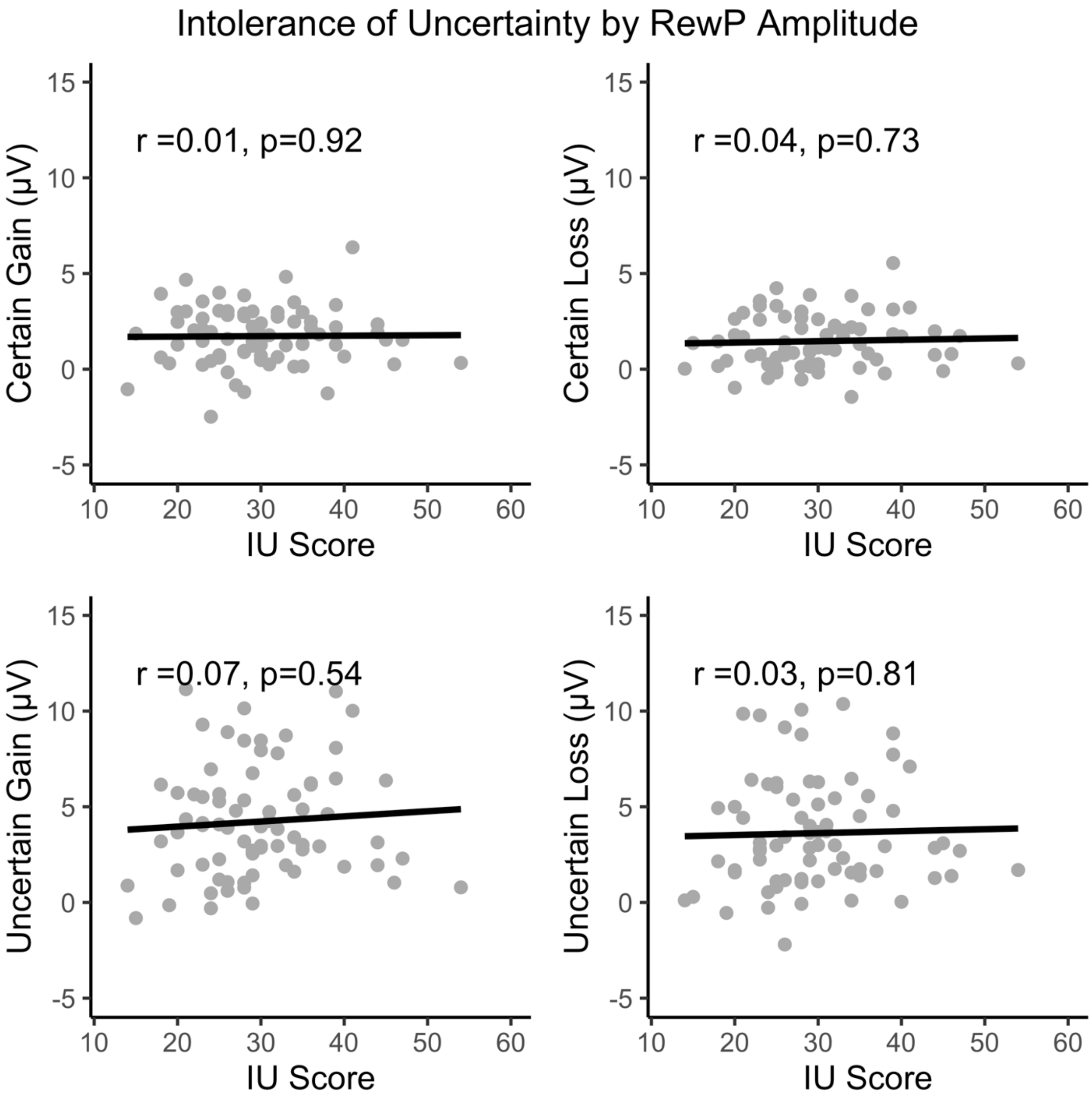


Figure S1: Correlations between intolerance of uncertainty total score and RewP amplitude for trial types out of the reward-cued task

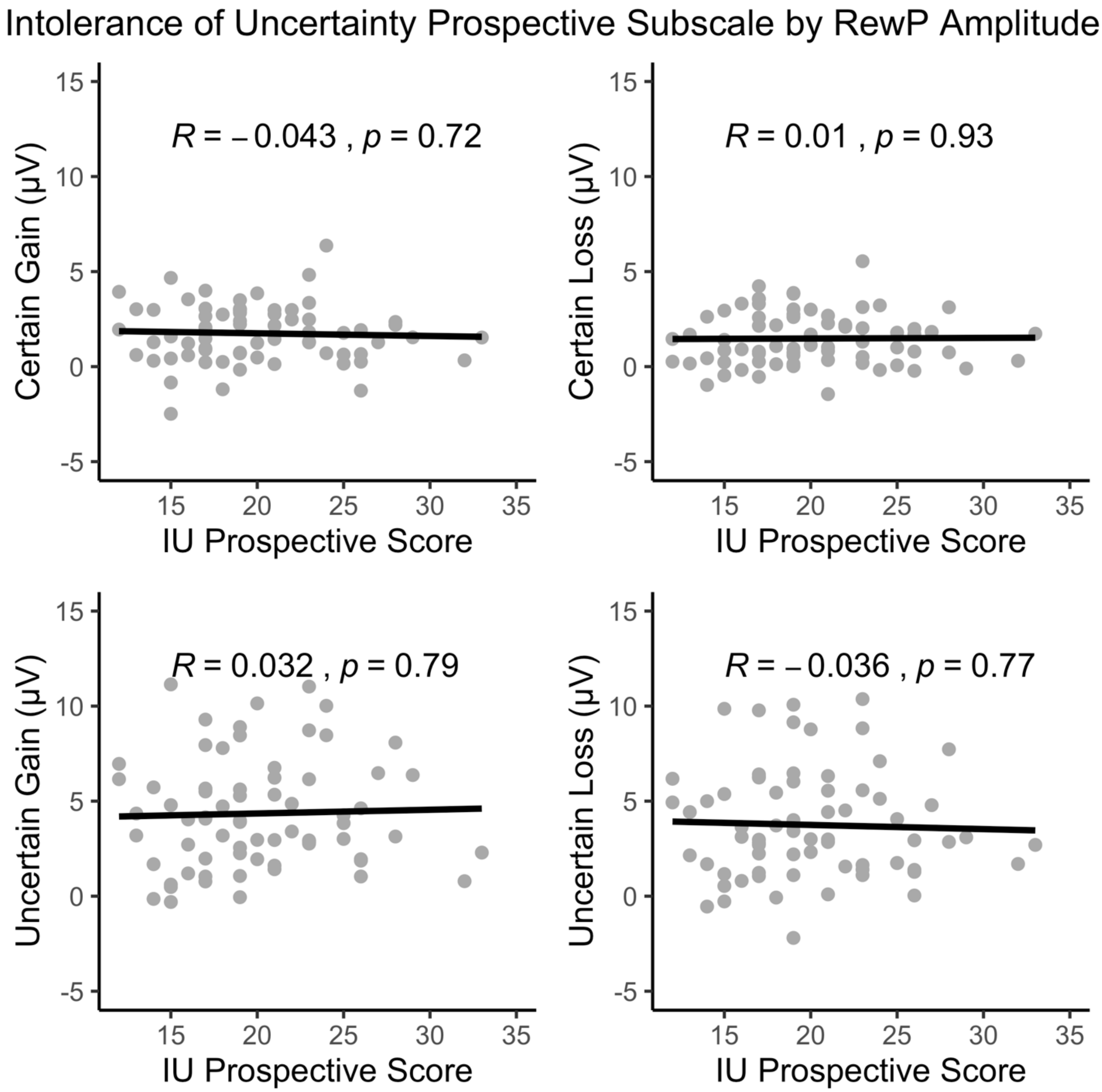


Figure S2: Correlations between intolerance of uncertainty prospective anxiety score and RewP amplitude for trial types out of the reward-cued task

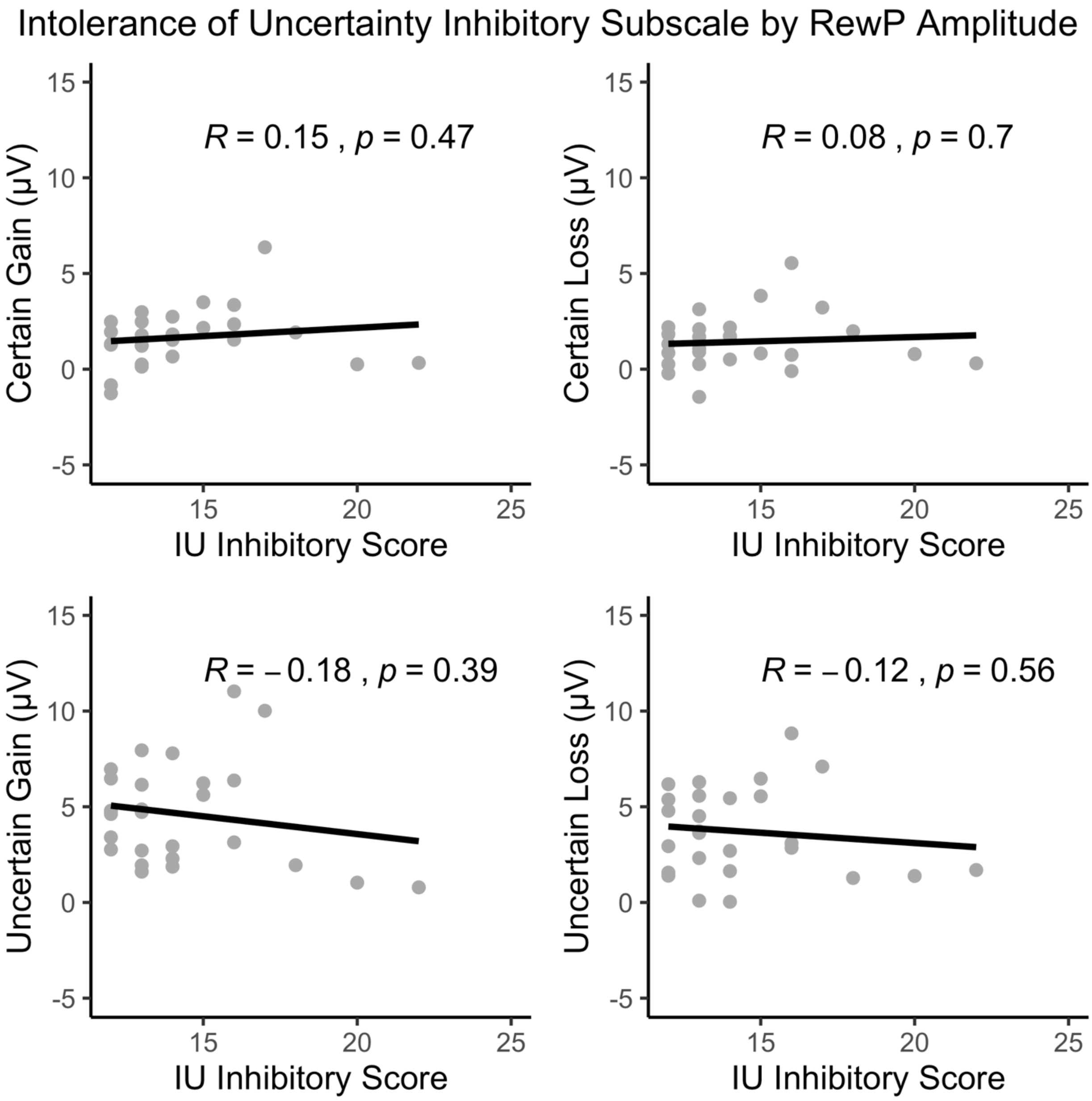


Figure S3: Correlations between intolerance of uncertainty inhibitory anxiety score and RewP amplitude for trial types out of the reward-cued task

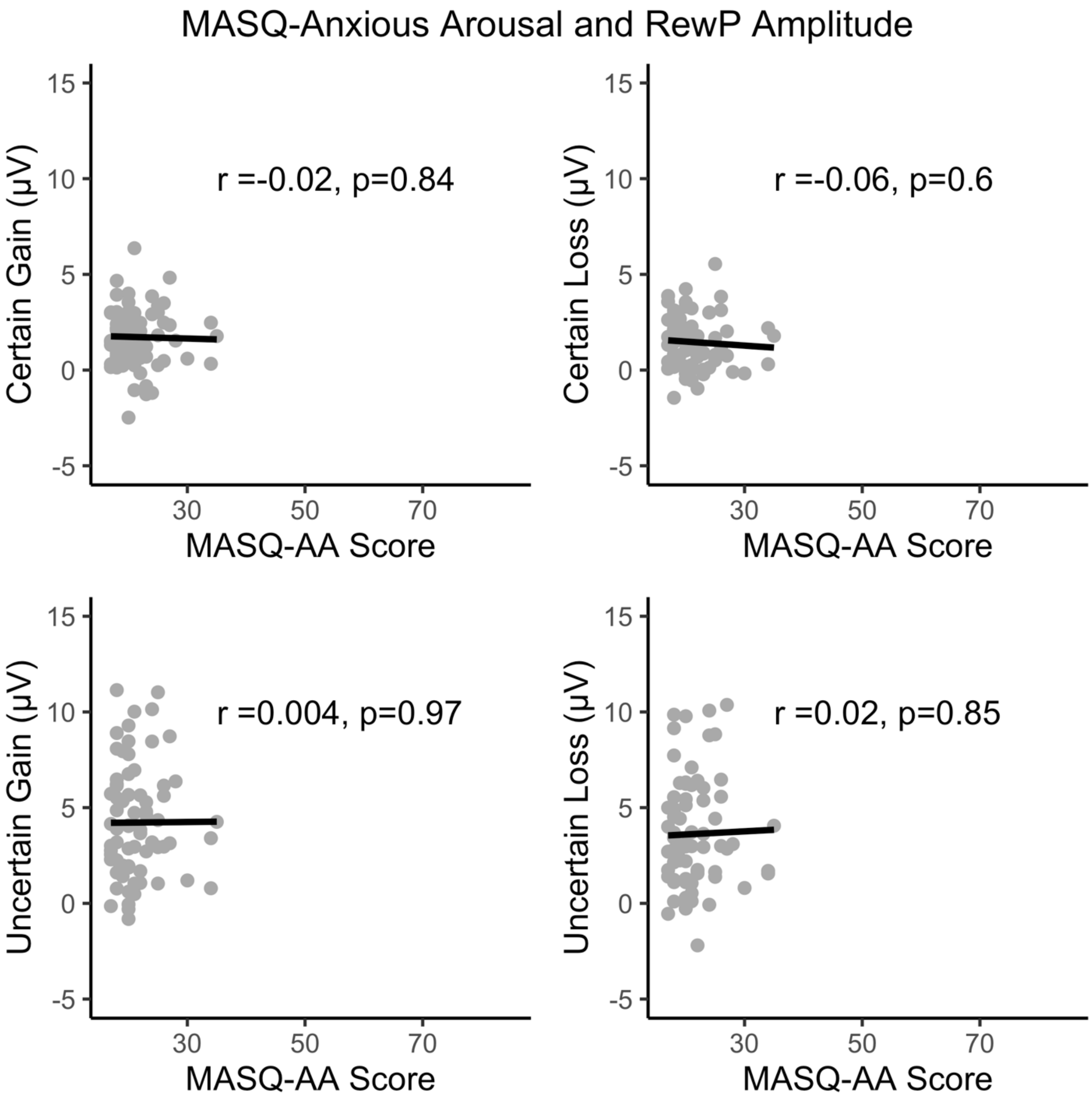


Figure S4: Correlations between MASQ-Anxious Arousal score and RewP amplitude for trial types out of the reward-cued task

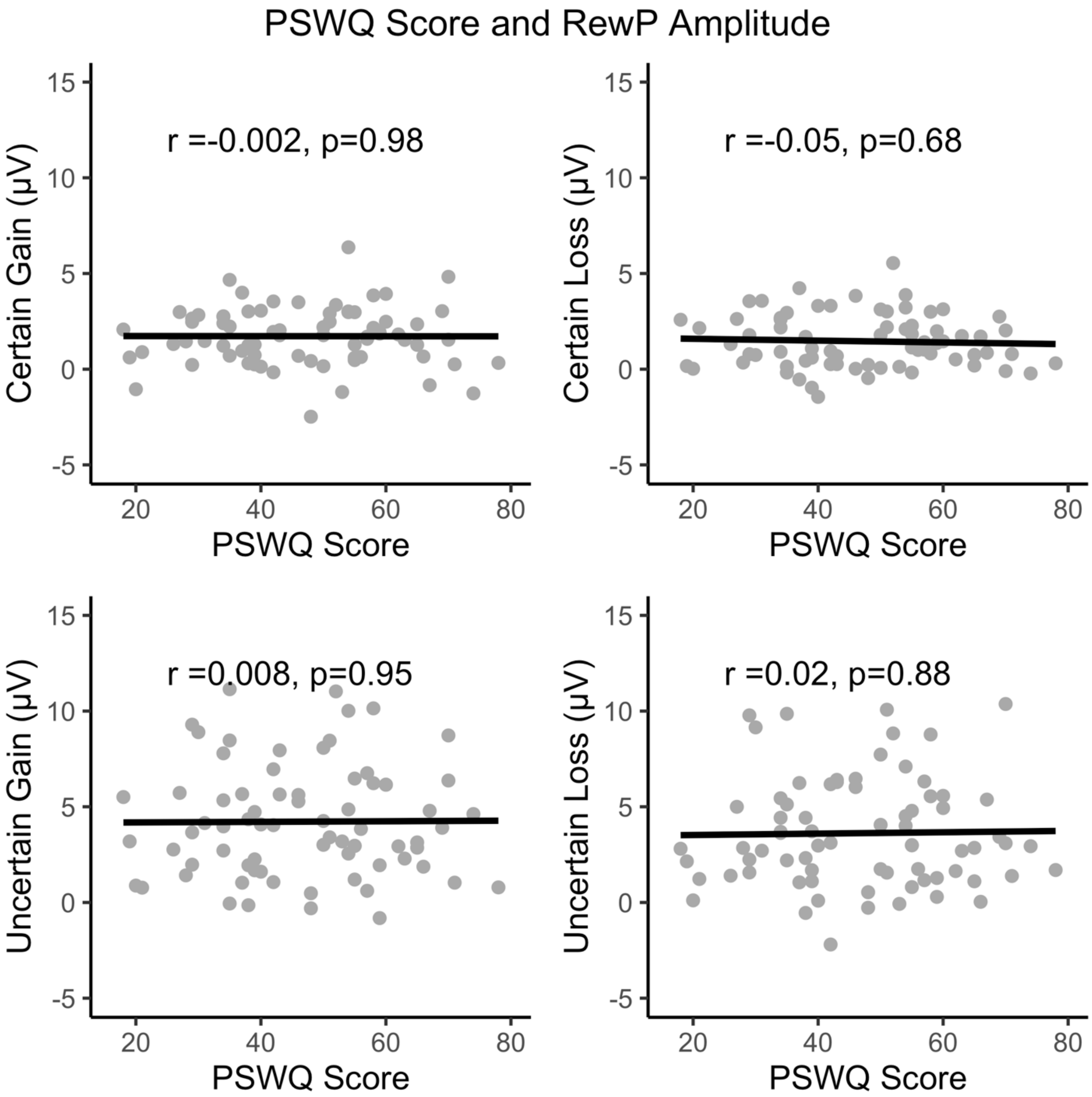


Figure S5: Correlations between Penn State Worry Questionnaire total scores and RewP amplitude for trial types out of the reward-cued task

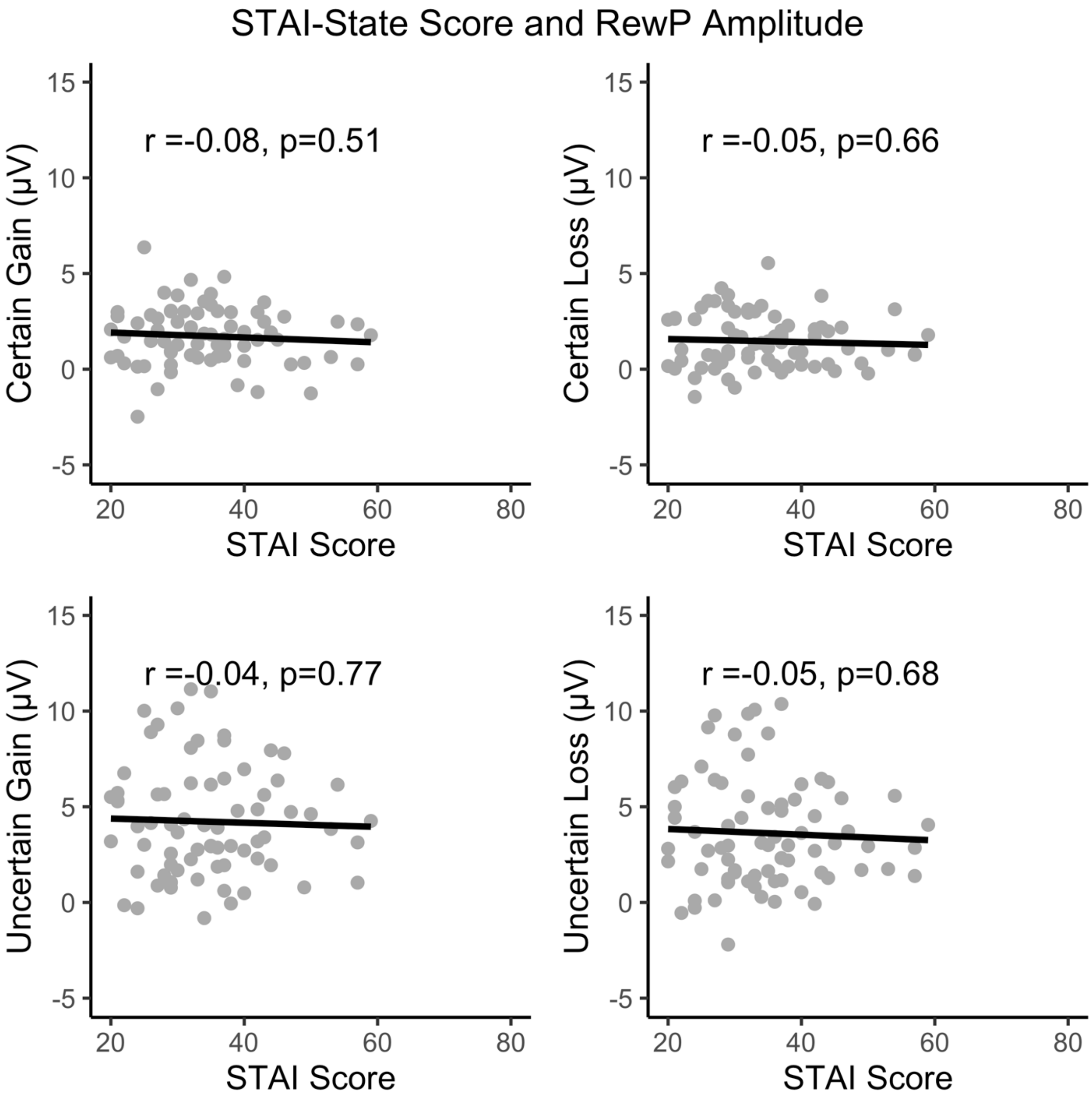


Figure S6: Correlations between STAI-State score and RewP amplitude for trial types out of the reward-cued task
